# Children are full of optimism, but those rose-tinted glasses are fading – reduced learning from negative outcomes drives hyperoptimism in children

**DOI:** 10.1101/2021.06.29.450349

**Authors:** Johanna Habicht, Aislinn Bowler, Madeleine E Moses-Payne, Tobias U Hauser

**Author notes:** **Corresponding author**, Tobias U. Hauser, Max Planck UCL Centre for Computational Psychiatry and Ageing Research, University College London, 10-12 Russell Square, London WC1B 5EH, United Kingdom, Phone: +44.

## Abstract

Believing that good things will happen in life is essential to maintain motivation and achieve highly ambitious goals. This optimism bias, the overestimation of positive outcomes, may be particularly important during childhood when motivation must be maintained in the face of negative outcomes. In a learning task, we have thus studied the mechanisms underlying the development of optimism bias. Investigating children (8-9 year-olds), early (12-13 year-olds) and late adolescents (16-17 year-olds), we find a consistent optimism bias across age groups. However, children were particularly hyperoptimistic, with the optimism bias decreasing with age. Using computational modelling, we show that this was driven by a reduced learning from worse-than-expected outcomes, and this reduced learning explains why children are hyperoptimistic. Our findings thus show that insensitivity to bad outcomes in childhood helps to prevent taking on an overly realistic perspective and maintain motivation.

## Introduction

Learning and knowing what is good for us is crucial for survival and a key part of development in human and non-human animals. Organisms are known to use trial-and-error learning to acquire and continuously adjust their knowledge and behaviour (Schultz et al., 1997). This reinforcement learning - the process of forming and adjusting expectations based on prediction errors (Sutton & Barto, 1998) - is known to converge to the optimal behaviour (Watkins & Dayan, 1992). This means that trial-and-error learning will allow the formation of correct and unbiased knowledge that maximises rewards (Körding & Wolpert, 2004; Sutton & Barto, 1998). Nevertheless, humans are known to be subject to many cognitive biases that distort these optimal representations (De Martino et al., 2006; Tversky & Kahneman, 1974).

A prominent distortion is an optimism bias, the tendency to see the world through rose-tinted glasses and to expect it to be better than reality actually is (Sharot, 2011). Even though this optimism bias may corrupt an adequate representation of one’s environment, it has been suggested to be beneficial because it boosts motivation and thus increases the likelihood of overcoming obstacles to achieve ambitious goals. Indeed, beneficial effects of optimism bias have been found in multiple domains, such as physical health (Rasmussen et al., 2009), mental health (Nolen-Hoeksema et al., 1992; Strunk et al., 2006) and professional development (Puri & Robinson, 2007).

The optimism bias might be of particular importance during development in childhood and adolescence. For example, an overly optimistic view of a dream job will allow a child to pursue their ambition and overcome the many obstacles along the way. In fact, the first studies investigating abstract concept knowledge revealed that children show features of optimism biases about the future in general (Bamford & Lagattuta, 2020; Fischer & Leitenberg, 1986), as well as their own future knowledge (Lockhart et al., 2017), and overestimate positive traits in themselves and others (Boseovski, 2010; Lockhart et al., 2002).

However, little is known about how this optimism bias arises mechanistically, especially in youths. A pioneering study in adults has investigated this process and has shown that an optimism bias indeed arises from a bias in learning, in which better-than-expected (positive) outcomes are weighted more strongly than worse-than-expected (negative) outcomes (Lefebvre et al., 2017). Using computational modelling, the authors showed that this optimism bias could be explained by an ‘optimistic learning’ bias, by means of a decreased learning rate for negative compared to positive prediction errors (Lefebvre et al., 2017).

Such an optimistic learning bias could be of particular importance when investigating optimism biases during development because a growing body of evidence indicates that prediction error related learning changes substantially during development (Cohen et al., 2020; Hauser et al., 2015; Nussenbaum & Hartley, 2019). In particular, several studies have suggested that overall learning from prediction errors increases during youth, although not ubiquitously so (for review cf (Nussenbaum & Hartley, 2019)). Whether a bias towards learning from positive compared to negative events exists during development and whether this optimistic learning bias leads to an optimism bias is unclear.

In this study, we investigate the development of optimism bias in childhood and adolescence, and how it is related to biases in reinforcement learning. We have chosen a sample that covers childhood (8-9 year-olds), early adolescence (12-13 year-olds) and late adolescence (16-17 year-olds) to understand how and when the optimism bias mechanistically changes over these key developmental stages. Using an ecologically realistic learning task (Hauser et al., 2017) that assessed youth’s ability to predict and learn from effortful attainment of reward, we show that children are hyperoptimistic with an increased optimism bias compared to early and late adolescents, and this bias is driven by a depleted learning rate for worse-than-expected outcomes.

## Methods

We recruited 108 participants from multiple schools across Greater London, UK. We only included participants who were in a certain age range (8-9 years old (yo), or 12-13 yo or 16-17 yo), and fluent in English. We allowed all children who provided consent to take part in the study but excluded from analysis those who had a history of neurological or psychiatric disorders, and visual impairment that was not corrected by the use of glasses or contact lenses. In total, nine participants were excluded from the analysis: 1 due to a pre-existing neurological condition (children group), 6 due to not understanding the task when asked questions about the task or not paying attention (as observed by experimenter; 2 from children group; 3 from early adolescents group; 1 from late adolescents group), 1 due to a technical problem (late adolescents group) and 1 due to low effort success rate (37.5%; children group). The final sample included 27 children (17 females; mean age = 9.32 ± 0.27), 38 early adolescents (20 females; mean age = 13.13 ± 0.31) and 34 late adolescents (21 females, mean age = 17.17 ± 0.28). All the behavioural findings reported were present when including those participants. These age ranges were selected to span late childhood, early- and late-adolescence. The groups did not differ in their age-adjusted IQ estimates (c.f. Table 1). We deliberately recruited participants from schools in socially diverse areas with lower socioeconomic status (SES) to counteract the currently overrepresented recruitment bias towards youth with higher SES (Fakkel et al., 2020). The sample size was selected assuming similar, medium to large effect sizes based on previous developmental studies and our own study using a similar task (Bamford & Lagattuta, 2020; Decker et al., 2015, 2016; Hauser et al., 2017; Lockhart et al., 2002, 2017). All participants provided written informed consent and participants under 16 provided written consent from a parent or legal guardian in addition to their own consent. Each participant was given a gift voucher of £7. The study was approved by the Research Ethics Committee of University College London (study number: 14261/001). Different tasks from the same participants are reported elsewhere (Bowler et al., 2021; Dubois et al., 2020; Moses-Payne et al., 2020). The current study was not preregistered. The data, analysis code and modelling toolbox are publicly available at https://github.com/DevComPsy/EL-development.

**Table 1.** Participants’ demographics. Age, gender and IQ scores for the three age groups. The groups did not differ in their IQ scores.

|  | ***Children***<br>***(8-9 years)*** | ***Early***<br>***adolescents***<br>***(12-13 years)*** | ***Late***<br>***adolescents***<br>***(17-18 years)*** |  |
| --- | --- | --- | --- | --- |
| ***Mean age***<br>***(±SD)*** | 9.32 ± 0.273 | 13.13 ± 0.312 | 17.17 ± 0.280 | |
| ***Gender (F/M)*** | 17 / 10 | 20 / 18 | 21 / 13 |  |
| ***IQ score***<br>***(WASI-II)*** | 93.81 ± 13.471 | 98.97 ± 13.663 | 97.91 ± 10.461 | F(2,96) = 1.377,<br>p=0.257 |

### Overview of procedure

The testing took place in a quiet room in the participants’ school in groups of three to four students. The adolescent groups completed the experiment in about 1.5 hours. For the youngest group the testing was spread over two sessions to reduce fatigue and took about 2 hours (including age-appropriate short breaks) in total to complete. Participants completed four tasks (other three reported elsewhere; e.g., (Bowler et al., 2021; Dubois et al., 2020; Moses-Payne et al., 2020)), a battery of questionnaires (used in conjunction with the other tasks) and short form of the WASI-II including the Vocabulary and Matrix Reasoning subtests (Wechsler, 1999) to estimate age-adjusted IQ. The participants were given both verbal and written instructions of the task before completing it. Children received longer instructions with examples to ensure that they understood the task. The order in which the tasks, questionnaires and WASI-II were administered, was pseudo-randomised across participants.

### Task

The goal of this study was to investigate how optimism bias and learning changed throughout development using an ecologically realistic effortful reward attainment task. We used a modified and child-friendly version of a previously established task (Hauser et al., 2017), where participants were helping an astronaut to fly a rocket to planets across the universe. They had to learn about the reward (1 to 7 gold coins) and an effort threshold that needed to be surpassed (amount of fuel the battery needed, which ranged between 42% and 92% of maximal button presses) in order to reach the planet to collect coins. Both reward magnitude and effort threshold slowly changed over time in a Gaussian random-walk–like manner. These trajectories were constructed so that reward and effort were decorrelated (cf Fig. 1). Importantly, the exerted effort was calibrated individually for each participant’s maximum number of presses. The maximum button presses were obtained in the practice session where the participants had to perform as many button presses as they could in 5 seconds. The calibration also included a staircase procedure, where the maximum effort was updated when the participant had exceeded their previous maximum effort in a trial.

**Figure 1.**
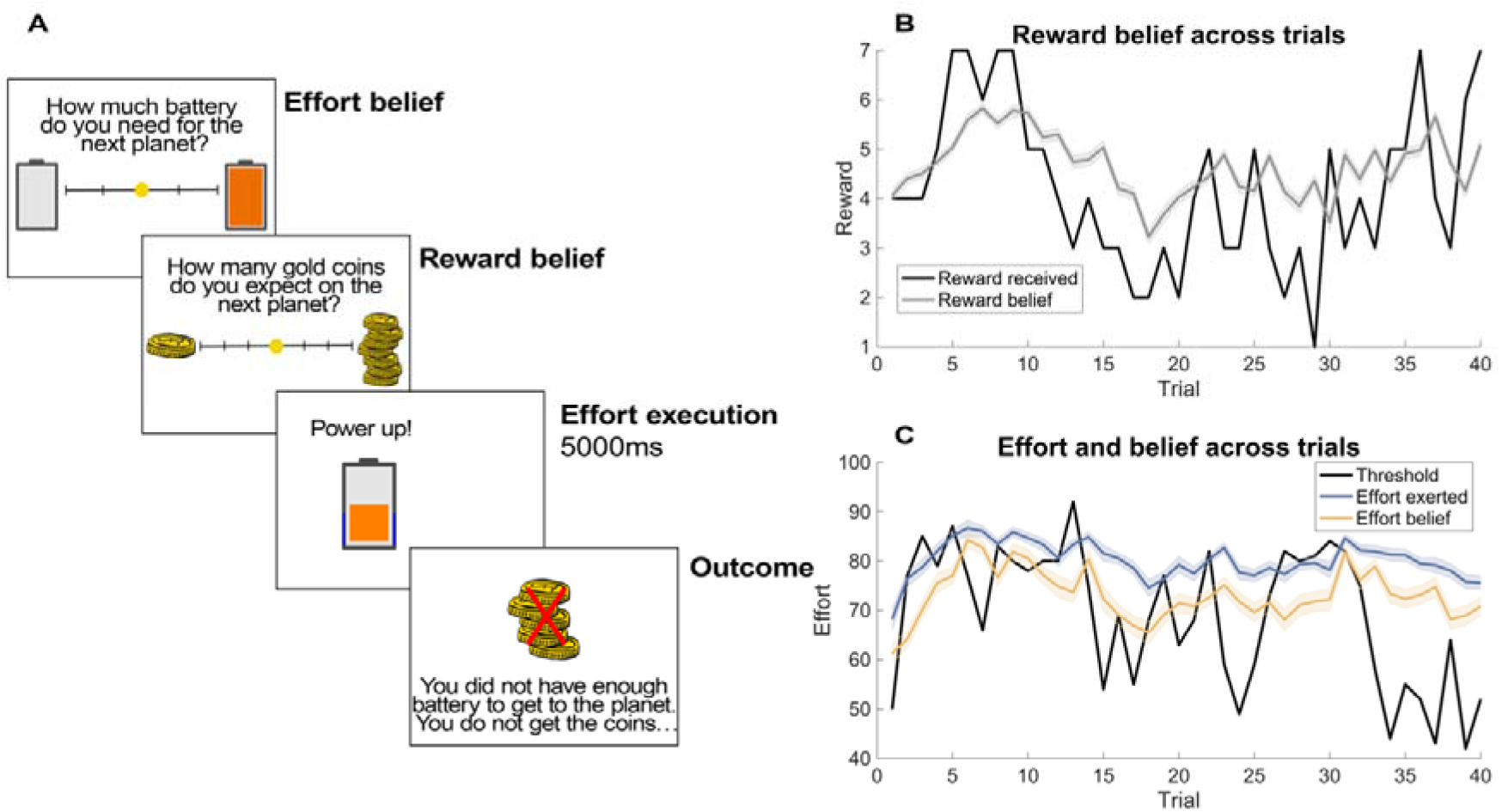
Task procedure and behaviour. (A) An effortful reward attainment learning task was used to assess learning and optimism bias. In the beginning of each trial, participants were asked to report their effort and reward belief. They then exerted the effort they believed was needed to obtain the rewards using button presses. The duration of button presses was kept constant across all trials to remove temporal discounting that may confound effort execution (Floresco et al., 2008). If successful, participants received coins that were revealed during outcome (here, 5 coins). If a participant exerted too little effort (i.e., did not exceed the necessary effort threshold), a cross was superimposed over the number of (potential) coins, indicating they will receive no reward on that trial, but still allowing them to learn about the potential reward. (B) Reward belief trajectory across trials (grey), compared to the actual reward received (black). (C) Trajectories of effort belief (yellow), effort exerted (blue) and the effort threshold (black) over trials. The line plots indicate the mean of beliefs and effort exerted ± 1 SEM.

In the beginning of each trial, the participants rated their belief about the amount of fuel needed to reach the next planet and their belief about how many coins they will get, ranging from 1 to 7. There was no time limit for reporting one’s beliefs. This was followed by 5 seconds of rapid manual button presses to fill up the battery. If the exerted effort was above the effort threshold, participants received the coins that were on display during outcome. If the participants’ effort did not exceed the threshold, a cross appeared above the number on display, which indicated that the participant did not receive any coins for that trial. Participants did not receive any explicit information about the threshold but had to learn from trial and error. After training, participants completed 40 trials.

### Multiple regression analyses

To assess the factors that influenced reward belief (Fig. 2C), we used multiple regression (fitglm function in Matlab) to predict the reward belief at each trial, respectively. As predictors, we entered the number of points (displayed during feedback) on the previous trial (previous reward; range: 1-7); and whether the force threshold was successfully surpassed on the previous trial (previous failure; failure coded as 1, success as −1). The last predictor was the participant’s reported reward belief on previous trial (previous reward belief; range: 1-7). The regression weights of the predictors were obtained for each individual and then tested for consistency across subjects using t-tests in a summary statistics approach.

**Figure 2.**
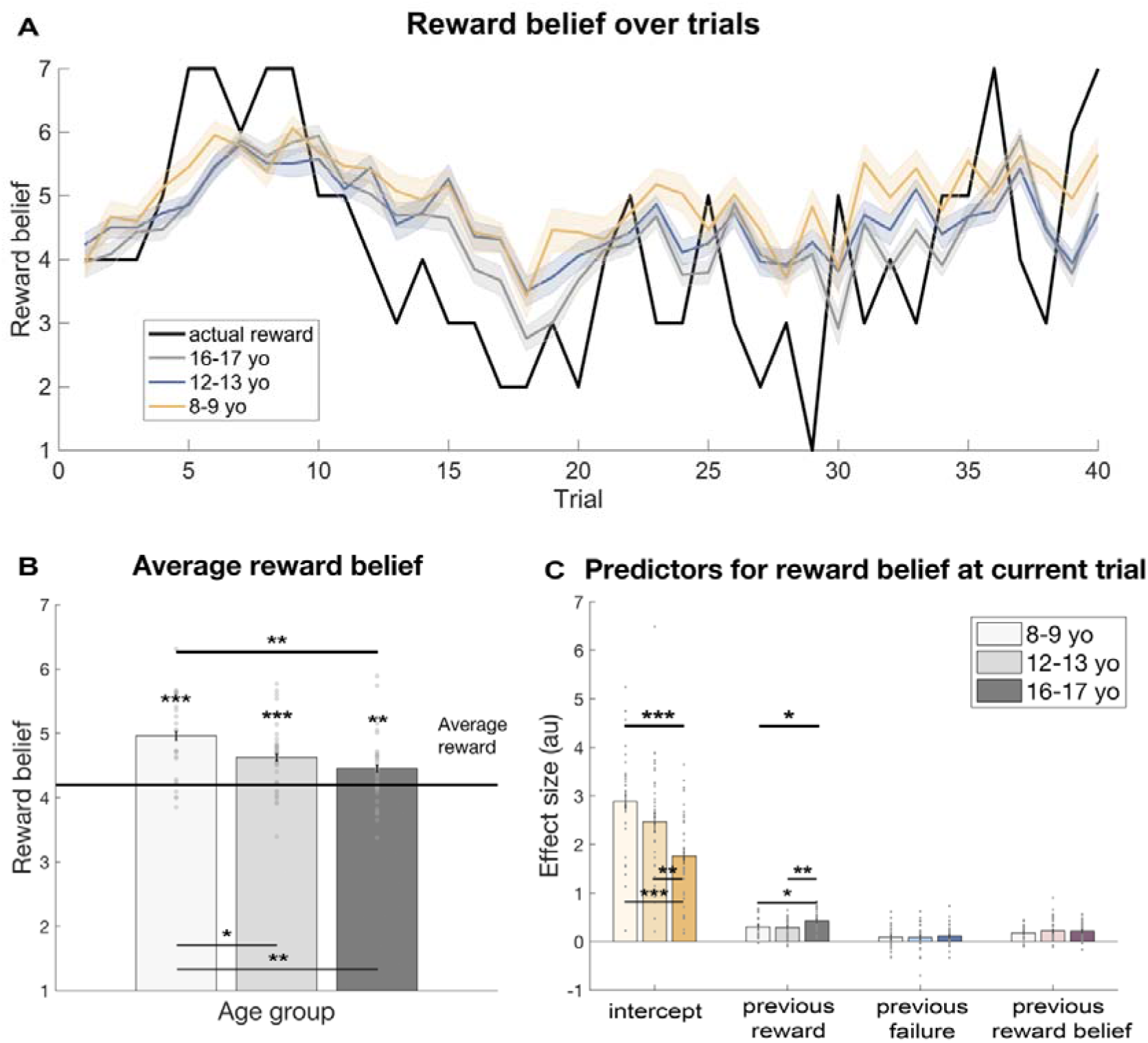
Hyperoptimism in youths decreases throughout development. (A) Reward belief trajectories split between the age groups over trials. Children had higher reward belief (yellow) throughout the task. The line plots indicate the reward belief on every trial averaged across participants ± 1 SEM. (B) An optimism bias (as measured by the average reward belief compared to the average reward in the task) was present in all groups. Importantly, this bias weakened during development. Children (8-9 year-olds) had a higher optimism bias compared to the adolescent groups (12-13 year-olds and 16-17 year-olds), revealing the emergence of more realistic expectations during adolescence. (C) Reward belief was predicted by the intercept, previous reward magnitude, failure to exceed the threshold on the previous trial, and participant’s own belief about previous reward magnitude. The intercept, which shows the general belief about reward magnitude over trials, was lower for late adolescents (16-17 year-olds) compared to children (8-9 year-olds) and early adolescents. Additionally, the previous reward magnitude predictor was higher for late adolescents compared to children and early adolescents, suggesting increased outcome sensitivity in late adolescence. Bar plots indicate mean ± 1 SEM; \*\*\**p* <.001; ** *p*<.01; \**p*<.05; yo, year-olds; au, arbitrary units.

### Statistical analyses

We compared behavioural measures using one-way ANOVAs with age group as between-subject factor (children, early adolescents, late adolescents). Significant effects were further explored using (independent samples) t-tests. We report effect sizes using partial eta squared (η^2^) for ANOVAs and Cohen’s d (d) for t-tests.

### Computational model

To investigate the computational mechanisms underlying age-related change in reward learning, we fitted six different variants of Rescorla-Wagner models (Rescorla & Wagner, 1972). We provide a summary of the models with key equations, parameter and model recovery in the Supplementary Material. Here we provide a brief description of the winning model and the key model parameters.

In this model, the subject starts with a prior belief μ_0_ about how big a reward will be, which is then adjusted based on the task feedback. This parameter can be seen as a static optimism bias, reflecting subjects’ prior expectation about how good a reward will be. Subsequently, the subject will learn from the task feedback by using two learning rates: if the outcome is better-than-expected (i.e. positive prediction error), the subject will update their belief using positive learning rate (α^+^), and when the outcome is worse-than-expected (i.e. negative prediction error), then the subject will update the belief using a negative learning rate (α^−^). If α^+^ > α^−^, then the subject learns more from positive outcomes compared to negative outcomes, in line with the precious account of optimistic learning bias (Lefebvre et al., 2017). The model also incorporates a noise parameter ξ that captures the noisiness of responding.

### Mediation analysis

We used mediation analysis to evaluate whether the effect of age on optimism bias was mediated by negative learning rate. We used standard notation to report mediation paths, where X represents the independent variable (age), Y represents the dependent variable (optimism bias) and M represents the mediating variable (negative learning rate). Relationship between X and M is expressed by path *a*, and relationship between M and Y is expressed by path *b*. The overall/total effect of X on Y is defined by path *c* and the direct effect of X on Y controlling for M is defined by path *c’*. The product *ab* defines the indirect effect of X on Y through M. If M mediates the relationship between X and Y, then the product *ab* should be significantly different from zero.

We used Mediation Toolbox in Matlab (https://github.com/canlab/MediationToolbox; Wager, Davidson, Hughes, Lindquist, & Ochsner, 2008; Wager et al., 2009) to perform the analysis. This toolbox is used to calculate mediation analysis based on a standard 3-variable path model (Baron & Kenny, 1986) with a bootstrap test for the statistical significance of the product *ab* (Efron & Tibshirani, 1993; Shrout & Bolger, 2002). This toolbox tests the significance of *ab* using the accelerated, bias-corrected bootstrap test (Efron & Tibshirani, 1993; Shrout & Bolger, 2002) with 10,000 bootstrap samples to test each of the *a*, *b* and *ab* path coefficients.

## Results

We examined the developmental differences in learning and optimism bias testing three groups of young people: 27 children (8-9 year-olds), 38 early adolescents (12-13 year-olds) and 34 late adolescents (16-17 year-olds). All subjects played a child-friendly, gamified version of a previously developed learning task (Hauser et al., 2017), in which subjects need to predict and learn from effortful attainment of reward. In essence, subjects needed to exert physical effort (button presses) to obtain a time-varying reward. Because both effort and reward changed over time (independently from each other), they had to constantly track and learn the changing effort demands and reward magnitudes through a process of trial and error (Fig. 1).

On every trial, participants were asked to report their beliefs about the reward they could obtain (number of gold coins, ranging from 1 to 7) and the effort needed to do so (battery fuel level) using a visual analogue scale (Fig. 1A). This allowed us to assess the subjects’ beliefs about both effort and reward. After reporting their belief, subjects needed to charge the battery of a space rocket to the level they believed was needed to reach the next planet quickly alternating between two button presses (physical effort exertion; fixed total charging time of 5 seconds). Subsequently, subjects were informed whether they exerted enough effort (i.e. reached the planet), and how many points they had won (or would have won). In the remainder of this paper, we will focus on reward learning and optimism bias, but detailed analyses on effort learning can be found in the supplementary material.

### Calibration ensures similar performance between groups

Before investigating optimism bias, we were interested in the overall performance of this task. We constructed the task so that actual performance was equal across all ages, thus preventing performance from confounding optimism bias. Concretely, we used a staircase procedure for each individual’s physical effort exertion to account for differences in physical or other abilities (cf Methods for details). This allowed the task to be challenging and achievable for everyone, whilst ensuring similar performance across groups. Indeed, we found that the number of successful trials (*F*(2,96) = 1.14, *p* = 0.323, *η^2^* = 0.023, Table S1), and the total points won in the task (*F*(2,96) = 1.33, *p* = 0.268, *η^2^* = 0.027) did not differ between the age groups. Furthermore, the average effort exerted (relative to the individual’s maximum effort) did not differ between groups (*F*(2,96) = 2.16, *p* = 0.121, *η* = 0.043), and neither did the variability of the effort exerted as measured by the effort standard deviation (*SD*; *F*(2,96) = 1.65, *p*=0.198, *η^2^* = 0.033). These findings thus allow us to compare optimism bias and reward learning between groups without having to account for biases in their performance.

### Hyperoptimism in children

Our key interest was to assess whether youths showed an optimism bias in this learning task. To this end, we compared the average reported reward belief across the entire task. We found that all groups consistently overestimated how many points they would get (Fig. 2B; all groups: *t*(98) = 77.77, *p* < 0.001; children: *t*(26) = 6.06, *p* < 0.001; early adolescents: *t*(37) = 5.04, *p* < 0.001; late adolescents: *t*(33) = 2.71, *p* = 0.010). This means that all participants’ expectations were higher than reality, which is the key characterisation of optimism bias (Sharot, 2011).

We next investigated whether this optimism bias changed with age showing a difference between age groups. We indeed found a significant age group effect (Fig. 2B, *F*(2,96) = 6.07, *p* = 0.003, *η^2^* = 0.112). Subsequent analyses showed that this was driven by children being most (hyper-)optimistic and that the extent of optimism bias decreased in adolescence (children vs early adolescents: *t*(63) = 2.29, *p* = 0.025, *d* = 0.565; children vs late adolescents: *t*(59) = 3.30, *p* = 0.002, *d* = 0.841; early vs late adolescents: *t*(70) = 1.27, *p* = 0.175, *d* = 0.323). This suggests that children are hyperoptimistic, and that this optimism bias is reduced but still present in adolescents. Furthermore, we found that children’s optimism bias increased over the course of the task, whereas optimism bias was more stable in both adolescent groups (cf Figure S2 for more details)

To make sure that children were not just more random in reporting their beliefs, we explored the variability of the participants’ reports and did not observe any age effects (SD of the reported reward belief; *F*(2, 96) = 1.68, *p* = 0.192, *η^2^* = 0.034; cf Figure S1 for reward belief distributions). Our findings thus suggest that an optimism bias is present in all youths, but it decreases as children get older.

### How are reward beliefs constructed?

To evaluate how participants constructed their reward beliefs, we performed a linear regression to predict reward belief at every trial. Given that the reward magnitudes were slowly drifting over time, the reward magnitude on the previous trial was a good indicator for the subsequent magnitude and is a reflection of a continuous learning process, such as reinforcement learning (Sutton & Barto, 1998). Assuming that participants followed a similar principle, we used the reward magnitude revealed on the previous trial (‘previous reward’) as a predictor, in addition to an intercept that captures the general belief of reward magnitude across the entire task. We found that both significantly predicted reward beliefs (previous reward: *M* = 0.34, *SD* = 0.219, *t*(98) = 15.39, *p* < 0.001; intercept: *M* = 2.30, *SD* = 1.741, t(98) = 13.14, *p* < 0.001).

We also evaluated whether further task aspects could influence the participants’ reward beliefs. In line with consistent reporting of one’s belief, we found that the current belief was also significantly related to the previous belief in an extended multiple regression (*M* = 0.20, *SD* = 0.197, *t*(98) = 10.29, *p* < 0.001). Lastly, we explored whether the effort had any influence on the reward beliefs. Indeed, we found some leakage from effort to reward belief formation, even though participants were instructed that these varied independently from each other. We found that previously exerted effort (i.e. button presses) did not predict the reward belief (*M* = 0.00, *SD* = 0.019, *t*(98) = 0.40, *p* = 0.692), but previous effort failure (i.e. whether one fulfilled the effort requirements on the previous trial) had a significant impact (*M* = 0.09, *SD* = 0.288, *t*(98) = 3.26, *p* = 0.002).

### Reward belief in late adolescents is more closely linked to previous reward

Next, we investigated whether there were any age differences in how the reward beliefs were constructed. We used the same linear regression with an intercept, previous reward, previous failure to exert enough effort, and one’s previous belief about reward as predictors, this time investigating age-dependent effects on these regressors. Comparing the regressors, we found significant age effect on the intercept (Fig. 2C; *F*(2,96) = 8.27, *p* < 0.001, *η^2^* = 0.147; children vs early adolescents: *t*(63) = 1.42, *p* = 0.161, *d* = 0.359; children vs late adolescents: *t*(59) = 4.26, *p* < 0.001, *d* = 1.085; early vs late adolescents: *t*(70) = 2.74, *p* = 0.008, *d* = 0.653). We also found that previous reward magnitude was integrated differently into reward belief in the different age groups (*F*(2,96) = 4.39, *p* = 0.015, *η*^2^ = 0.084). This was mainly driven by late adolescents adjusting their reward belief more according to previous reward compared to early adolescents (*t*(70) = −2.83, *p* = 0.006, *d* = 0.668) and children (*t*(59) = −2.27, *p* = 0.027, *d* = 0.580). There were no differences between children and early adolescents (*t*(63) = 0.235, *p* = 0.815, *d* = 0.059). As the previous reward magnitude is an indicative measure for learning, it suggests that learning may differ between age groups.

We did not find any differences in how one’s own previous belief about reward was used to construct current reward belief (*F*(2,96) = 0.40, *p* = 0.674, *η*^2^ = 0.008), suggesting no age-related differences in how participants integrate their own internal reward belief into their learning. There were also no differences in how previous failure impacted on reward belief construction (*F*(2,96) = 0.16, *p* = 0.850, *η^2^* = 0.003). Our findings thus indicate that the difference in optimism was driven, at least in part, by altered learning. However, this analysis is insensitive to more subtle learning biases, such as differences in learning for positive and negative outcomes.

### Computational mechanisms underlying reward belief

To better understand the mechanisms underlying the developmental effects on learning and optimism bias, we developed a computational model similar to the previous study reporting learning mechanisms in optimism bias (Lefebvre et al., 2017). In short (cf supplementary material for detailed model description), our model learns about the reward magnitude using a modified Rescorla-Wagner (Rescorla & Wagner, 1972) prediction error learning rule. In this model, learning is driven by a prediction error, the difference between expected and received reward. This prediction error is then used to update a reward belief using a learning rate α. Similar to the previous work on optimism bias, we used two separate learning rates, one for positive (α^+^) and one for negative (α^−^) prediction errors. If α^+^ > α^−^, one learns more from better-than-expected than worse-than-expected outcomes (optimistic learning). If α^−^ > α^+^, one deploys pessimistic learning. If α^+^ = α^−^, one would learn equivalently from positive and negative events (realistic learning). This is based on the assumption that optimistic learning would lead to optimism bias over time (Lefebvre et al., 2017). In addition, our model incorporated a prior in reward belief (μ_0_) as an additional free parameter, capturing any reward bias that was present at the outset of the task. Lastly, we used a noise parameter ξ to capture the noisiness of responding.

We fitted the model to each participant’s behaviour (cf Supplementary material for detailed model comparison and fitting) and found that all groups showed optimistic learning as α^+^ was higher than α^−^ for all ages (paired t-tests; children: *t*(26) = 4.04, *p* < 0.001, early adolescents: *t*(37) = 4.63, *p* < 0.001, late adolescents: *t*(33) = 2.25, *p* = 0.032). Moreover, the difference between positive and negative learning rate (α^+^ - α^−^) was highly correlated with the optimism bias (average reward belief) in our task (*r* = 0.582, *p* < 0.001, partial correlation controlling for age: *r* = 0.557, *p* < 0.001), suggesting that the optimistic learning bias is associated with the optimism bias.

### Negative learning rate increases with age

To understand how the model parameters changed across age to drive the effect on optimism bias, we compared the parameters across age groups. We found significant age effect on the negative learning rate α^−^ (Fig. 3A; *F*(2,96) = 8.34, *p* < 0.001, *η^2^* = 0.148), but no effect on the positive learning rate α^+^ (Fig. 3B; *F*(2,96) = 0.09, *p* = 0.914, *η^2^* = 0.002). When further investigating α^−^, we found the age difference was driven by children learning less from negative prediction errors compared to adolescents (children vs early adolescents: *t*(54) = −3.66, *p* = 0.001, *d* = 0.868; children vs late adolescents: *t*(37.25) = −4.03, *p* < 0.001, *d* = 0.988; early vs late adolescents: *t*(48.46) = −1.91, *p* = 0.063, *d* = 0.461). This means that while positive learning remains stable across ages, negative learning increases substantially, diminishing the optimistic learning bias in the older participants.

**Figure 3.**
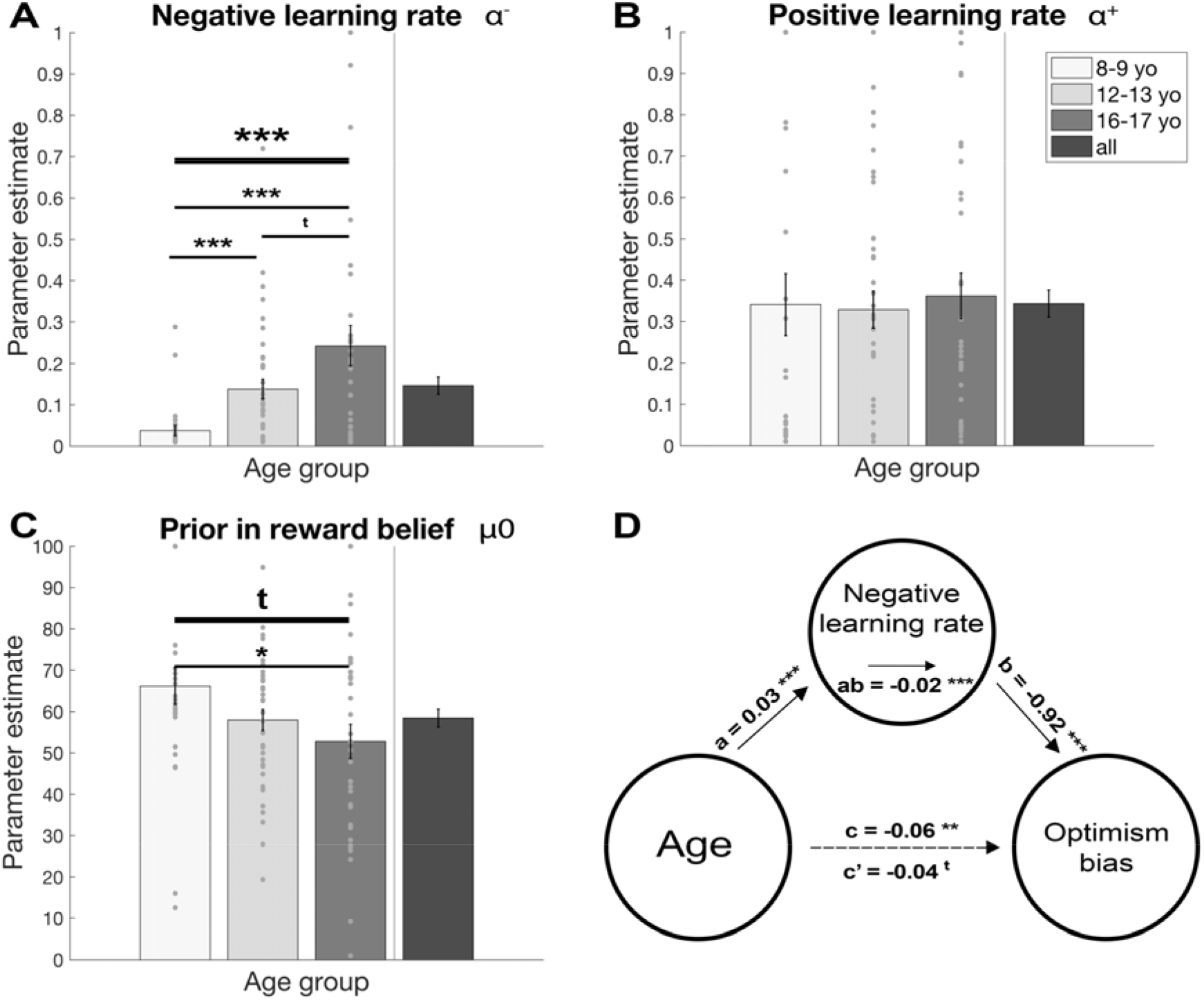
Model parameters reveal differences in optimistic learning. The parameters of the bestfitting model reveal that children (8-9 yo) differed from the adolescent groups (12-13 yo and 16-17 yo) in their negative learning rate. (A) Children have a lower negative learning rate compared to both adolescent groups, whereas the positive learning rate is similar across all age groups (B). This means that children show a strong optimism learning bias, which diminishes with age. Additionally, children have marginally higher prior in reward belief compared to late adolescents (C). The relationship between age and optimism bias was mediated by negative learning rate (D). Mean values are shown, the *c* path represents the total effect of age on optimism bias and *c’* represents the direct effect of age on optimism bias when controlling for the negative learning rate. The *a* path represents the effect on age on negative learning rate and *b* path represents the effect on negative learning rate on optimism bias. The *ab* path is the indirect effect of age on optimism bias through negative learning rate. Our findings suggest that the age-related decrease in optimism bias was primarily driven by the increase in the negative learning rate. \*\*\**p* < 0.001; ** *p* < 0.01; \**p* < 0.05; t *p* < 0.10; yo, year-olds.

We further explored the other task parameters and found a marginally significant age effect on μ_0_ (Fig., 3C; *F*(2,96) = 3.00, *p* = 0.054, *η^2^* = 0.059). This was driven by children having higher prior beliefs about rewards compared to late adolescents (children vs early adolescents: *t*(63) = 1.69, *p* = 0.097, *d* = 0.410; children vs late adolescents: *t*(59) = 2.17, *p* = 0.034, *d* = 0.560; early vs late adolescents: *t*(55.52) = −1.05, *p* = 0.297, *d* = 0.252). This suggests that besides an age-dependent learning optimism bias, there is also a trend towards a prior optimism bias in children.

### Increase in negative learning rates mediates age-related decline in optimism bias

To assess whether the observed optimism bias was driven by the changes in learning that we observed (especially the negative learning rates), we conducted a mediation analysis testing whether the negative learning rate mediated the association between age and optimism bias. Our mediation analysis confirmed the significant associations between age and negative learning rate (path a: *M* = 0.03, *SE* = 0.01, *z* = 4.54, *p* < 0.001) and between age and optimism bias (path c: *M* = −0.06, *SE* = 0.02, *z* = −3.28, *p* = 0.001). Moreover, we found a significant association between negative learning rate and optimism bias (path b: *M* = −0.92, *SE* = 0.22, *z* = −3.93, *p* < 0.001).

Our mediation analysis further revealed that the association between age and optimism bias was mediated by negative learning rate (path ab: *M* = −0.02, *SE* = 0.01, *z* = −3.44, *p* < 0.001), and the association between age and optimism bias was substantially diminished and remained only marginally significant when accounting for negative learning rate (path c’: *M* = −0.04, *SE* = 0.02, *z* = −1.96, *p* = 0.051). Importantly, none of the other model parameters mediated the decrease in optimism bias (results not shown). These results suggest the decrease in optimism bias is mediated by the increased learning from events where the outcome is worse than expected, reducing the optimism learning bias.

## Discussion

Studying optimism bias in the context of a learning task revealed the mechanisms underlying the changes in optimism bias from childhood and adolescence. Our findings provide novel evidence that optimism bias exists at a more general reinforcement learning level during development, and the extent of optimism bias decreases from childhood to adolescence. Furthermore, we show that this optimism bias is associated with learning less from worse-than-expected outcomes, and the change in learning from negative prediction errors mediates the age effect on optimism bias. In other words, as children become older, they learn more from negative outcomes, lose their hyperoptimism, and become more realistic.

We use individuals’ reported beliefs about the future reward as an insight to their optimism bias (Stankevicius et al., 2014) and show that children and adolescents exhibit an optimism bias about rewards. To our knowledge, this is the first study that examines optimism bias in trial-and-error learning across development rather than optimism bias in more abstract contexts, such as future vulnerability (Moutsiana et al., 2013) or future knowledge (Lockhart et al., 2017). The difference between the previous studies (Bamford & Lagattuta, 2020; Fischer & Leitenberg, 1986; Lockhart et al., 2017; Moutsiana et al., 2013) and ours is that previous studies investigated (hypothetical) future events, whereas our study allowed to trace learning and the emergence of optimism bias instantly. It is interesting that the optimism bias is still present when rewards are immediate and experienced as in the current study rather than imagined in the future. Our findings demonstrate that children show hyperoptimism compared to both early and late adolescents, and thus extend the previous accounts that assessed optimism bias in abstract contexts in children, such as aspects of the future (Bamford & Lagattuta, 2020; Fischer & Leitenberg, 1986; Lockhart, Chang, & Story, 2002). By investigating development across childhood, early and late adolescence, our findings further extend previous studies comparing adults to children (Lockhart, Chang and Story, 2002; Lockhart, Goddu and Keil, 2017) and provide a more fine-grained developmental resolution.

We use computational modelling to show that the observed optimism bias arises from optimistic learning, i.e. learning less from negative than positive prediction errors. This aligns with pioneering research in adults showing that more optimistic individuals disregard negative information about the future (Sharot et al., 2011), and they learn less from worse-than-expected than better-than-expected outcomes in reinforcement learning (Lefebvre et al., 2017). However, it is important to consider that optimistic learning, and consequently optimism bias, can be task-specific, as the adaptability of asymmetric learning (degree of positive and negative learning rates in relation to each other) depends on the reward distribution (Cazé & Van Der Meer, 2013) and perceived controllability of the task (Chambon et al., 2020; A. O. Cohen et al., 2020; Dorfman et al., 2019). Thus, the optimism bias that arises from optimistic learning may be adaptive to task environments.

Our findings suggest that learning from negative outcomes increases during development, whereas learning from positive outcomes stays constant during childhood and adolescence. This is in line with previous research, in which younger age was associated with diminishing negative information about how likely an individual is to encounter adverse life events (e.g. home burglary) (Moutsiana et al., 2013), but age did not affect how one updated their beliefs about positive information. Yet, our results are different from other previous studies that found a decrease in negative learning rates with age (Hauser et al., 2015; Rodriguez Buritica et al., 2018; Van Den Bos et al., 2012). However, two of the three studies compared either children to adults (Rodriguez Buritica et al., 2018), or adolescents to adults (Hauser et al., 2015), which do not necessarily contradict with our findings, as the change in negative learning rate might have an inverse u-shaped trajectory, peaking during adolescence (J. R. Cohen et al., 2010). Moreover, inconsistent findings about learning rates may reflect the task-specific adaptations that are dependent on the reward statistics of the task environment and how optimal learning is characterised in the task (Nussenbaum & Hartley, 2019). In the other mentioned studies (Hauser et al., 2015; Rodriguez Buritica et al., 2018; Van Den Bos et al., 2012), the statistical properties of the task environment differ from our task, and thus the extent to which one should weigh the recent outcomes to make the most optimal decisions differs between tasks. In other words, every task has a different optimal strategy to learn depending on the task environment (Cazé & Van Der Meer, 2013; Dorfman et al., 2019), thus making learning rates task-specific and incomparable between different task environments. Indeed, Nussenbaum and Hartley (2019) suggest that the inconsistencies in how the learning rates change with age between tasks suggest that there may be a change how individuals adapt their behaviour with age – individuals may become better with age at adapting their behaviour to make the most optimal decisions in a particular environment.

Interestingly, the increase in the negative learning rate seems to be at the heart of the age-dependent change in optimism bias. In our mediation analysis, we find that the increase in negative learning rate mediates the decrease of optimism bias from childhood to adolescence. No similar effect was found when investigating whether the other computational parameters mediate this association. Our findings thus suggest that the change in learning from worse-than-expected outcomes is the mechanism underlying the hyperoptimism found in children: as children get older, they incorporate more negative information from the environment and thus become more realistic. It would be interesting to further assess this mechanism across the lifespan to understand how optimism bias changes later in life.

It is important to consider other factors that may have influenced learning and optimism bias in our task. Recent papers have shown that the optimism biases may change based on the perceived controllability of the task (Dorfman et al., 2019). Concretely, a higher perceived controllability by the individual has been shown to be associated with higher optimism biases (Klein & Helweg-Larsen, 2002). Contextual beliefs, such as whether the external environment is good or bad, may influence the extent of learning from negative and positive outcomes (Dorfman et al., 2019). In our study, we did not experimentally modify the controllability or the valence of the task, and can therefore not draw any conclusions about how these processes change with age, or how they depend on learning. Interestingly, a very recent study by Cohen and colleagues assessed the influence of locus of control on optimism bias in the context of development. In their study, the authors found that children did not use their beliefs about the causal structure of the environment to guide learning in the same manner as adolescents and adults (Cohen et al., 2020). It would thus be interesting to further assess whether these were two distinct processes that matured during adolescence, or whether they were driven by the same mechanisms. Secondly, it is noteworthy that we had only outcomes in the positive valence domain in this task (i.e. no potential losses were included). It is thus interesting to conjecture whether losses would differently affect the optimism bias. Some previous studies that have investigated optimism bias (e.g Moutsiana et al., 2013; Sharot et al., 2011; Sharot, 2011), have indeed used bad/negative valence outcomes such as negative life events, as the worst outcomes. Interestingly, Lefebreve and colleagues have compared different task versions that either had reward omission as the worst outcome (positive valence domain), or had losses as worst outcomes (negative valence). When investigating the optimism bias in adults, the authors found that this difference in reward valence did not affect the optimistic learning bias (Lefebvre et al., 2017). However, it would be interesting to investigate the influence of reward omission versus bad outcome on optimism bias during development. Another limitation of our study was not preregistering it, which we would carefully consider in our next studies.

The increased learning from negative prediction errors with age, and the subsequent decrease in optimism bias, may be related to the development of dopaminergic system during adolescence (Blakemore & Robbins, 2012), but the exact, likely complex, mechanisms remain unknown (Hauser et al., 2019). Dopamine has been previously shown to reduce learning from negative information (Sharot et al., 2012), and in our previous study in adults using a similar task (Hauser et al., 2017) we found that these reward prediction errors (PEs) were encoded in dopamine-rich areas, such as the ventral striatum (VS). Moreover, activity in VS has been shown to be associated with learning from negative PEs (Lefebvre et al., 2017), and sensitivity to rewarding outcomes in VS has been revealed to peak during adolescence in an inverted U-shaped function (Braams et al., 2015; Hauser et al., 2015; Somerville et al., 2010; Somerville & Casey, 2010; Walker et al., 2017). Furthermore, the activity in response to negative compared to positive feedback was shown to increase during development (Peters & Crone, 2017). In sum, this suggests that the decrease in hyperoptimism throughout childhood and adolescence might be related to dopamine functioning and its effect on learning from negative prediction errors, but needs to be investigated in more detail.

It is fascinating to speculate why this over-optimism in children has developed. It may be a protective mechanism that helps children to maintain motivation and encourage perseverance (Boseovski, 2010). With an optimistic view on future outcomes, children may be more likely to pursue trial-and-error approaches (Boseovski, 2010), explore more (Dubois et al., 2020) and be more open-minded (Bjorklund & Green, 1992; Lucas et al., 2014), which is beneficial for learning in the long run. Furthermore, the optimism bias may encourage optimal learning strategies that lead to higher performance in specific environments, such as tasks with low reward probabilities (Cazé & Van Der Meer, 2013) or environments where one has high controllability over their actions (Chambon et al., 2020; Dorfman et al., 2019). Optimism may also act as a protective mechanism for the development of psychiatric disorders, such as depression. Lack of optimism bias has been shown to be higher in individuals with depression (Strunk et al., 2006), driven by increased learning from negative events (Garrett et al., 2014). Given that early adolescence is a period of heightened risk for mental health conditions (Kessler et al., 2005) it would be interesting to study how trajectories of learning imbalances predict the risk of psychiatric symptoms. Thus, future studies could map the individual developmental trajectories of psychiatric symptoms, learning and optimism bias with age in longitudinal studies.

In summary, we show that children are hyperoptimistic, and that this optimism bias decreases throughout adolescence. Using computational modelling, we show that the reduction in learning from negative prediction errors drives this age-related decrease in hyperoptimism. This optimism bias might act as a driving factor for children’s resilience and motivate them to keep on trying different approaches to learn more in the long run.

## Conflict of interest

The authors declare no competing financial interests.

## Acknowledgements

We thank the children, adolescents, their schools (especially Sydney Russell School, Baden Powell School and Holmleigh Primary School) and families for taking part in our study. We thank Shona Waters for her help with collecting the data. TUH is supported by a Wellcome Sir Henry Dale Fellowship (211155/Z/18/Z), a grant from the Jacobs Foundation (2017-1261-04), the Medical Research Foundation, a 2018 NARSAD Young Investigator grant (27023) from the Brain & Behavior Research Foundation, and an ERC Starting Grant. The Max Planck UCL Centre is a joint initiative supported by UCL and the Max Planck Society. The Wellcome Centre for Human Neuroimaging is supported by core funding from the Wellcome Trust (203147/Z/16/Z). A CC BY or equivalent licence is applied to the Author Accepted Manuscript arising from this submission, in accordance with the grant’s open access conditions.

## Notes

### Competing Interest Statement

The authors have declared no competing interest.

## References

Bamford, C., & Lagattuta, K. H. (2020). Optimism and Wishful Thinking: Consistency Across Populations in Children’s Expectations for the Future. Child Development. https://doi.org/10.1111/cdev.13293

Baron, R. M., & Kenny, D. A. (1986). The moderator-mediator variable distinction in social psychological research: Conceptual, strategic, and statistical considerations. Journal of Personality and Social Psychology. https://doi.org/10.1037//0022-3514.51.6.1173

Bjorklund, D. F., & Green, B. L. (1992). The adaptive nature of cognitive immaturity. American Psychologist. https://doi.org/10.1037/0003-066X.47.1.46

Blakemore, S. J., & Robbins, T. W. (2012). Decision-making in the adolescent brain. In Nature Neuroscience. https://doi.org/10.1038/nn.3177

Boseovski, J. J. (2010). Evidence for “Rose-colored glasses”: An examination of the positivity bias in young children’s personality judgments. Child Development Perspectives. https://doi.org/10.1111/j.1750-8606.2010.00149.x

Bowler, A., Habicht, J., Moses-Payne, M. E., Steinbeis, N., Moutoussis, M., & Hauser, T. U. (2021). Children perform extensive information gathering when it is not costly. Cognition, 208, 104535. https://doi.org/10.1016/j.cognition.2020.104535

Braams, B. R., van Duijvenvoorde, A. C. K., Peper, J. S., & Crone, E. A. (2015). Longitudinal changes in adolescent risk-taking: A comprehensive study of neural responses to rewards, pubertal development, and risk-taking behavior. Journal of Neuroscience. https://doi.org/10.1523/JNEUROSCI.4764-14.2015

Cazé, R. D., & Van Der Meer, M. A. A. (2013). Adaptive properties of differential learning rates for positive and negative outcomes. Biological Cybernetics. https://doi.org/10.1007/s00422-013-0571-5

Chambon, V., Théro, H., Vidal, M., Vandendriessche, H., Haggard, P., & Palminteri, S. (2020). Information about action outcomes differentially affects learning from self-determined versus imposed choices. Nature Human Behaviour. https://doi.org/10.1038/s41562-020-0919-5

Cohen, A. O., Nussenbaum, K., Dorfman, H. M., Gershman, S. J., & Hartley, C. A. (2020). The rational use of causal inference to guide reinforcement learning strengthens with age. Npj Science of Learning. https://doi.org/10.1038/s41539-020-00075-3

Cohen, J. R., Asarnow, R. F., Sabb, F. W., Bilder, R. M., Bookheimer, S. Y., Knowlton, B. J., & Poldrack, R. A. (2010). A unique adolescent response to reward prediction errors. Nature Neuroscience. https://doi.org/10.1038/nn.2558

De Martino, B., Kumaran, D., Seymour, B., & Dolan, R. J. (2006). Frames, biases and rational decision-making in the human brain. Science. https://doi.org/10.1126/science.1128356

Decker, J. H., Lourenco, F. S., Doll, B. B., & Hartley, C. A. (2015). Experiential reward learning outweighs instruction prior to adulthood. Cognitive, Affective and Behavioral Neuroscience. https://doi.org/10.3758/s13415-014-0332-5

Decker, J. H., Otto, A. R., Daw, N. D., & Hartley, C. A. (2016). From Creatures of Habit to Goal-Directed Learners: Tracking the Developmental Emergence of Model-Based Reinforcement Learning. Psychological Science. https://doi.org/10.1177/0956797616639301

Dorfman, H. M., Bhui, R., Hughes, B. L., & Gershman, S. J. (2019). Causal Inference About Good and Bad Outcomes. Psychological Science. https://doi.org/10.1177/0956797619828724

Dubois, M., Bowler, A., Moses-Payne, M. E., Habicht, J., Steinbeis, N., & Hauser, T. U. (2020). Tabula-rasa exploration decreases during youth and is linked to ADHD symptoms. BioRxiv, 2020.06.11.146019. https://doi.org/10.1101/2020.06.11.146019

Efron, B., & Tibshirani, R. J. (1993). An Introduction to the Bootstrap. In An Introduction to the Bootstrap. https://doi.org/10.1007/978-1-4899-4541-9

Fakkel, M., Peeters, M., Lugtig, P., Zondervan-Zwijnenburg, M. A. J., Blok, E., White, T., van der Meulen, M., Kevenaar, S. T., Willemsen, G., Bartels, M., Boomsma, D. I., Schmengler, H., Branje, S., & Vollebergh, W. A. M. (2020). Testing sampling bias in estimates of adolescent social competence and behavioral control. Developmental Cognitive Neuroscience. https://doi.org/10.1016/j.dcn.2020.100872

Fischer, M., & Leitenberg, H. (1986). Optimism and Pessimism in Elementary School-Aged Children. Child Development. https://doi.org/10.2307/1130655

Floresco, S. B., Tse, M. T. L., & Ghods-Sharifi, S. (2008). Dopaminergic and glutamatergic regulation of effort- and delay-based decision making. Neuropsychopharmacology. https://doi.org/10.1038/sj.npp.1301565

Garrett, N., Sharot, T., Faulkner, P., Korn, C. W., Roiser, J. P., & Dolan, R. J. (2014). Losing the rose tinted glasses: Neural substrates of unbiased belief updating in depression. Frontiers in Human Neuroscience. https://doi.org/10.3389/fnhum.2014.00639

Hauser, T. U., Eldar, E., & Dolan, R. J. (2017). Separate mesocortical and mesolimbic pathways encode effort and reward learning signals. Proceedings of the National Academy of Sciences of the United States of America. https://doi.org/10.1073/pnas.1705643114

Hauser, T. U., Iannaccone, R., Walitza, S., Brandeis, D., & Brem, S. (2015). Cognitive flexibility in adolescence: Neural and behavioral mechanisms of reward prediction error processing in adaptive decision making during development. NeuroImage. https://doi.org/10.1016/j.neuroimage.2014.09.018

Hauser, T. U., Will, G. J., Dubois, M., & Dolan, R. J. (2019). Annual Research Review: Developmental computational psychiatry. Journal of Child Psychology and Psychiatry and Allied Disciplines. https://doi.org/10.1111/jcpp.12964

Kessler, R. C., Berglund, P., Demler, O., Jin, R., Merikangas, K. R., & Walters, E. E. (2005). Lifetime prevalence and age-of-onset distributions of DSM-IV disorders in the national comorbidity survey replication. In Archives of General Psychiatry. https://doi.org/10.1001/archpsyc.62.6.593

Klein, C. T. F., & Helweg-Larsen, M. (2002). Perceived control and the optimistic bias: A meta-analytic review. In Psychology and Health. https://doi.org/10.1080/0887044022000004920

Körding, K. P., & Wolpert, D. M. (2004). Bayesian integration in sensorimotor learning. Nature. https://doi.org/10.1038/nature02169

Lefebvre, G., Lebreton, M., Meyniel, F., Bourgeois-Gironde, S., & Palminteri, S. (2017). Behavioural and neural characterization of optimistic reinforcement learning. Nature Human Behaviour. https://doi.org/10.1038/s41562-017-0067

Lockhart, K. L., Chang, B., & Story, T. (2002). Young children’s beliefs about the stability of traits: Protective optimism? Child Development. https://doi.org/10.1111/1467-8624.00480

Lockhart, K. L., Goddu, M. K., & Keil, F. C. (2017). Overoptimism about future knowledge: Early arrogance? Journal of Positive Psychology. https://doi.org/10.1080/17439760.2016.1167939

Lucas, C. G., Bridgers, S., Griffiths, T. L., & Gopnik, A. (2014). When children are better (or at least more open-minded) learners than adults: Developmental differences in learning the forms of causal relationships. Cognition. https://doi.org/10.1016/j.cognition.2013.12.010

Moses-Payne, M., Habicht, J., Bowler, A., Steinbeis, N., & Hauser, T. (2020). I know better! Emerging metacognition allows adolescents to ignore false advice. PsyArXiv. https://doi.org/10.31234/osf.io/gb9f4

Moutsiana, C., Garrett, N., Clarke, R. C., Lotto, R. B., Blakemore, S. J., & Sharot, T. (2013). Human development of the ability to learn from bad news. Proceedings of the National Academy of Sciences of the United States of America. https://doi.org/10.1073/pnas.1305631110

Nolen-Hoeksema, S., Girgus, J. S., & Seligman, M. E. P. (1992). Predictors and Consequences of Childhood Depressive Symptoms: A 5-Year Longitudinal Study. Journal of Abnormal Psychology. https://doi.org/10.1037/0021-843X.101.3.405

Nussenbaum, K., & Hartley, C. A. (2019). Reinforcement learning across development: What insights can we draw from a decade of research? In Developmental Cognitive Neuroscience. https://doi.org/10.1016/j.dcn.2019.100733

Peters, S., & Crone, E. A. (2017). Increased striatal activity in adolescence benefits learning. Nature Communications. https://doi.org/10.1038/s41467-017-02174-z

Puri, M., & Robinson, D. T. (2007). Optimism and economic choice. Journal of Financial Economics. https://doi.org/10.1016/j.jfineco.2006.09.003

Rasmussen, H. N., Scheier, M. F., & Greenhouse, J. B. (2009). Optimism and physical health: A meta-analytic review. Annals of Behavioral Medicine. https://doi.org/10.1007/s12160-009-9111-x

Rescorla, R. A., & Wagner, A. R. (1972). A theory of Pavlovian conditioning: variations in the effectiveness of reinforcement and nonreinforcement. In A. H. Black & W. F. Prokasy (Eds.), Classical Conditioning II: Current Research and Theory. Appleton Century-Crofts.

Rodriguez Buritica, J. M., Heekeren, H. R., Li, S. C., & Eppinger, B. (2018). Developmental differences in the neural dynamics of observational learning. Neuropsychologia. https://doi.org/10.1016/j.neuropsychologia.2018.07.022

Schultz, W., Dayan, P., & Montague, P. R. (1997). A Neural Substrate of Prediction and Reward. Science, 275(5306), 1593 LP – 1599. https://doi.org/10.1126/science.275.5306.1593

Sharot, T. (2011). The optimism bias. In Current Biology. https://doi.org/10.1016/j.cub.2011.10.030

Sharot, T., Guitart-Masip, M., Korn, C. W., Chowdhury, R., & Dolan, R. J. (2012). How dopamine enhances an optimism bias in humans. Current Biology. https://doi.org/10.1016/j.cub.2012.05.053

Sharot, T., Korn, C. W., & Dolan, R. J. (2011). How unrealistic optimism is maintained in the face of reality. Nature Neuroscience. https://doi.org/10.1038/nn.2949

Shrout, P. E., & Bolger, N. (2002). Mediation in experimental and nonexperimental studies: New procedures and recommendations. Psychological Methods. https://doi.org/10.1037/1082-989X.7.4.422

Somerville, L. H., & Casey, B. J. (2010). Developmental neurobiology of cognitive control and motivational systems. In Current Opinion in Neurobiology. https://doi.org/10.1016/j.conb.2010.01.006

Somerville, L. H., Jones, R. M., & Casey, B. J. (2010). A time of change: Behavioral and neural correlates of adolescent sensitivity to appetitive and aversive environmental cues. In Brain and Cognition. https://doi.org/10.1016/j.bandc.2009.07.003

Stankevicius, A., Huys, Q. J. M., Kalra, A., & Seriès, P. (2014). Optimism as a Prior Belief about the Probability of Future Reward. PLoS Computational Biology. https://doi.org/10.1371/journal.pcbi.1003605

Strunk, D. R., Lopez, H., & DeRubeis, R. J. (2006). Depressive symptoms are associated with unrealistic negative predictions of future life events. Behaviour Research and Therapy. https://doi.org/10.1016/j.brat.2005.07.001

Sutton, R. S., & Barto, A. G. (1998). Reinforcement Learning: An Introduction. IEEE Transactions on Neural Networks. https://doi.org/10.1109/tnn.1998.712192

Tversky, A., & Kahneman, D. (1974). Judgment under uncertainty: Heuristics and biases. Science. https://doi.org/10.1126/science.185.4157.1124

Van Den Bos, W., Cohen, M. X., Kahnt, T., & Crone, E. A. (2012). Striatum-medial prefrontal cortex connectivity predicts developmental changes in reinforcement learning. Cerebral Cortex. https://doi.org/10.1093/cercor/bhr198

Wager, T. D., Davidson, M. L., Hughes, B. L., Lindquist, M. A., & Ochsner, K. N. (2008). Prefrontal-Subcortical Pathways Mediating Successful Emotion Regulation. Neuron. https://doi.org/10.1016/j.neuron.2008.09.006

Wager, T. D., Waugh, C. E., Lindquist, M., Noll, D. C., Fredrickson, B. L., & Taylor, S. F. (2009). Brain mediators of cardiovascular responses to social threat. NeuroImage. https://doi.org/10.1016/j.neuroimage.2009.05.043

Walker, D. M., Bell, M. R., Flores, C., Gulley, J. M., Willing, J., & Paul, M. J. (2017). Adolescence and reward: Making sense of neural and behavioral changes amid the chaos. In Journal of Neuroscience. https://doi.org/10.1523/JNEUROSCI.1834-17.2017

Watkins, C. J. C. H., & Dayan, P. (1992). Q-learning. Machine Learning. https://doi.org/10.1007/bf00992698

Wechsler, D. (1999). Wechsler Abbreviated Scale of Intelligence. The Psychological Corporation: Harcourt Brace & Company.

