## Supplementary Material for "Children are full of optimism, but those rose-tinted glasses are fading – reduced learning from negative outcomes drives hyperoptimism in children"

**Performance in groups**

| | **Children (8-9 years)** | **Early adolescents (12-13 years)** | **Late adolescents (17-18 years)** |
|---|---|---|---|
| Proportion of successful trials (%) | 76.30 (±10.524) | 76.12 (±10.146) | 73.09 (±8.462) |
| Total points won | 130.89 (±17.328) | 129.74 (±16.810) | 124.71 (±14.438) |
| Effort exerted out of max effort (%) | 81.21 (±4.986) | 80.58 (±4.800) | 78.97 (±3.454) |
| Exerted effort variability | 13.75 (±4.055) | 12.37 (±4.451) | 11.80 (±4.101) |

**Table S1.** The performance variables in groups. The values are reported as mean (±SD).

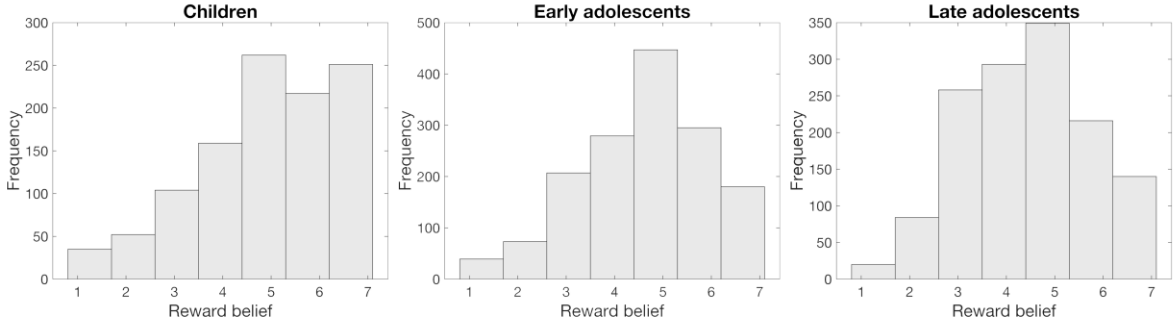


**Figure S1.** Participants reward belief distributions across age groups. Participants reward belief reports were well distributed across the scale, showing a similar pattern of responses across age groups, expect for children having higher frequency of reporting reward belief of 6 and 7.

**Optimism bias**

To investigate whether optimism bias changed over the course of the task (as a indirect approximation of learning), we examined optimism bias for each block separately. We found a marginally significant increase in optimism bias from block 1 to block 2 in children (paired t-test: t(26) = -1.96, p = 0.061, d = 0.377). We did not observe this effect neither in early adolescents (t(37) = 0.24, p = 0.813, d = 0.039), nor in late adolescents (t(33) = -0.32, p = 0.751, d = 0.055). This is in line with our main finding that over the course of the task, children become more optimistic because they learn less from negative outcomes, and that this effect is much less reduced in adolescents.

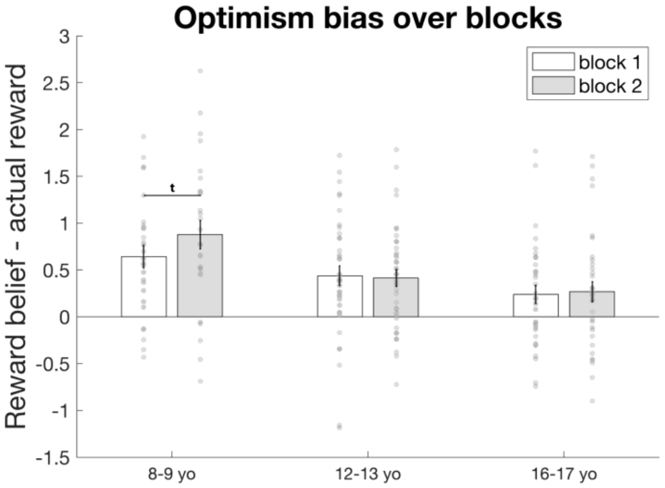


**Figure S2.** Participants' optimism bias over blocks for each age group. The optimism bias is calculated as the the difference between the individual's average reward belief in a block and the average actual reward received in that block. Early and late adolescents had a stable optimism bias over the two blocks, whereas in children the optimism bias increased (marginally) in the second block.

### Computational Modelling

To capture reward learning in this task, we compared different variants of a Rescorla-Wagner model (Rescorla & Wagner, 1972) fitted to subjects' reward ratings, similar to previous modelling approaches capturing an optimism (learning) bias (Lefebvre, Lebreton, Meyniel, Bourgeois-Gironde, & Palminteri, 2017).

We assumed that subjects started with a prior belief $\mu_0$ about how big a reward will be, which is then adjusted based on the task feedback. Model selection (Fig. S3) revealed that models that had a free $\mu_0$ parameter outperformed models that kept it as a fixed parameter (set at 50; free parameter range: 0-100). This parameter can be seen as a static optimism bias, reflecting subjects' prior expectation about how good a reward will be.

Subjects' belief about the reward they would get was then updated throughout the game using a prediction error δ, which was the difference between the reward belief $\mu_t$ and the reward they received at this trial $r_t$

$$\delta_t = \mathrm{r_t} - \mu_\mathrm{t}$$

This predictions error was then used to update the belief using one or multiple learning rates α (range: 0-1) for both positive and negative prediction errors

$$\mu_{t+1} = \mu_t + \alpha\delta_t$$

In the simpler (worse fitting) model family, a single learning rate was used for all outcomes. Alternatively, and in line with previous modelling of optimism bias (Lefebvre et al., 2017) we used two separate learning rates, one for positive ($\alpha^+$) and one for negative prediction errors

($\alpha^{-}$). A higher learning rate for positive prediction errors was assumed to be an indication of optimistic learning.

$$\mu_{t+1} = \mu_t + \alpha^{+}\delta_t \qquad if \ \ \delta > 0$$

$$\mu_{t+1} = \mu_t + \alpha^{-}\delta_t \qquad if \ \ \delta < 0$$

As the subjects used a continuous rating scale, rather than a simple (binary) choice, we assumed that the belief was transformed into a rating (i.e. choice) using a Gaussian with mean $\mu_t$ and standard deviation 10 (fixed parameter).

$$\pi_{rating} = \mathcal{N}(\mu_t, 10)$$

We additionally considered noise in the reporting using a 'noise floor' parameter $\xi$, which increased the likelihood to report the rating anywhere along the rating scale (range: 0-.1).

$$\pi_{rating} = \mathcal{N}(\mu_t, 10) + \xi$$

The $\pi_{rating}$ was normalised so that the policy summed up to 1

$$\pi_{rating} = \frac{\pi_{rating}}{\sum_{1-100} \pi_{rating}}$$

**Model fitting**

We fitted six different models to each subject's behaviour using maximum likelihood estimation, estimating the parameters using a genetic algorithm as implemented in Matlab (R2020a). Model selection was performed using summed BIC (Schwarz, 1978) across all subjects. For illustration, BIC was also presented for each group separately, revealing similar results in each group (cf Figure S3).

**Model parameter recovery analysis**

To assess how precise the model parameters of interest were, we ran a parameter recovery analysis for the winning model. We simulated 500 agents with parameter values randomly drawn from the respective parameter ranges, and then fit the simulated task behaviour in the same way as the real participants (cf Model fitting). To assess the fit between simulated and fitted parameters, we used Pearson correlations.

We found that the three parameters of interest all had good to excellent parameter recoverability: positive learning rate $r_{\alpha^+} = .87$, negative learning rate $r_{\alpha^-} = .86$, prior reward belief $r_{\mu_0} = .59$ (all *p*s < 0.001). Only the noise floor parameter ξ performed poorly (r = -.04) at the given large value range that we used for model fitting to avoid boundary problems (see Fig S4 for empirical values that have a much lower than range). Given that models with this noise still improve model fits (Fig. S3), this suggest that adding some noise improves the model, but that the degree of noise is of limited interpretability. In sum this means that our parameters-of-interest are well recoverable, but that the noise floor parameter ξ should not be interpreted.

**Model recovery analysis**

We were further interested how well that the two best models (1α vs 2α model, both with $\mu_0$ and ξ) were recoverable. We thus simulated 500 agents for each of the models (randomly drawing parameter values as described in parameter recovery analysis), and then fitted them with both models. Recoverability analysis was then done based on the BIC values, in accordance with the model selection procedure.

We found that 86% if the agents simulated with the 1α model were correctly attributed to the correct generative model (i.e. had the lowest BIC for the 1α model), and that also 82% of the 2α agents were correctly attributed. This means that these models were well-dissociable in our task.

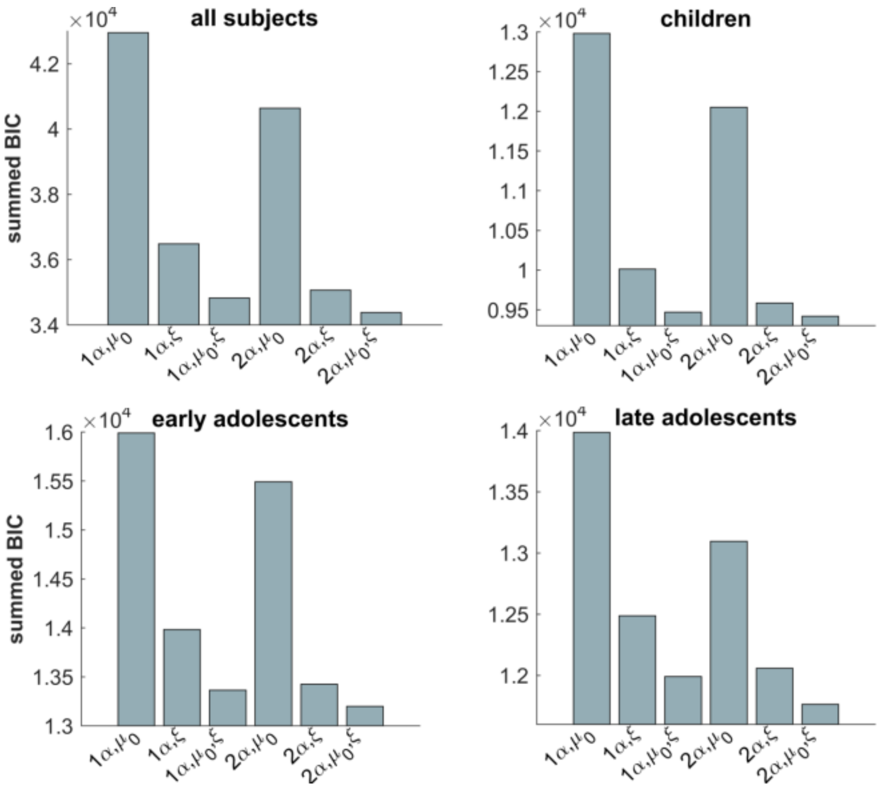


**Figure S3.** Model selection results. We used summed BIC to compare between six competing models (across all subjects, top left). Model comparison revealed that the best fitting model was the one that had separate learning rates (2α) for positive and negative prediction errors. In addition, this model had a free parameter for the prior reward expectation ($\mu_0$) and a noise response parameter (ξ).

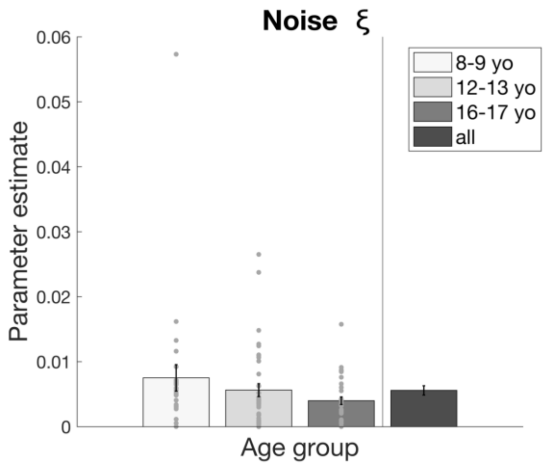


**Figure S4.** The noise parameter (ξ) of the best fitting model is similar in all groups. yo, year-olds.

### Effort analysis

*No group differences in bias about effort threshold*

To assess whether there was a bias in how the effort threshold was perceived, we compared the average effort belief to the average threshold. Importantly, participants did not have any explicit feedback on the threshold but rather had to learn from trial and error. We found that in all groups the average effort belief was higher than the threshold (children: $t(26) = 2.72$, $p = 0.011$, early adolescents: $t(37) = 2.62$, $p = 0.013$, late adolescents: $t(33) = 3.11$, $p = 0.003$). However, as participants did not receive any explicit information about the threshold, it is evident that they assumed the threshold to be higher as they had to surpass it to succeed. We did not find any differences between age groups how the effort belief was perceived ($F(2,96) = 0.27$, $p = 0.767$, $\eta2 = 0.006$) or in the SD of the submitted beliefs ($F(2,96) = 1.84$, $p = 0.164$, $\eta2 = 0.037$).

Next, we assessed whether effort belief or exerted effort were associated with reward belief. We found a weak correlation between the exerted effort and reward belief weak ($M = 0.21$, $SD= 0.211$, $t(98) = 9.97$, $p < 0.001$), as well as a weak correlation between effort belief and reward belief ($M = 0.35$, $SD = 0.266$, $t(98) = 13.23$, $p < 0.001$).

*Effort belief construction in all groups*

To assess how effort belief was constructed explicitly, we performed a multiple regression analysis that predicted effort belief on every trial. We found that previous effort (Fig S5B; $M = 0.10$, $SD = 0.202$, $t(98) = 5.04$, $p < 0.001$), previous failure ($M = 0.37$, $SD = 0.252$, $t(98) = 14.71$, $p < 0.001$), previous reward ($M = 0.14$, $SD = 0.193$, $t(98) = 7.14$, $p < 0.001$) and previous effort belief ($M = 0.31$, $SD = 0.248$, $t(98) = 12.31$, $p < 0.01$) all contributed to estimating effort belief on current trial. The previous effort and failure regressors highlight that participants took their own physical effort exertion, as well as whether they exerted too little effort and failed, into account to compute their beliefs about the threshold. Furthermore, they used previous reward information to compute their effort belief. This can be considered as a 'bleed-in' effect as the reward and effort are independent, thus participants should not use reward feedback to construct their explicit beliefs about effort. Lastly, participants took their own previous effort belief into account, suggesting that their belief construction was not random, but was built on their previous beliefs.

*Effort belief construction develops over childhood*

We found that there were age differences in how previous effort was included in effort belief (Fig S5B; $F(2,96) = 5.82$, $p = 0.004$, $\eta 2 = 0.108$), driven by difference between children and adolescents (children vs early adolescents: $t(63) = -2.71$, $p = 0.009$, $d = 0.674$ ; children vs late adolescents: $t(59) = -3.14$, $p = 0.003$ $d = 0.801$; early vs late adolescents: $t(70) = 0.60$, $p = 0.552$, $d = 0.140$). In addition, there were age differences how the previous failure affected the effort belief ($F(2,96) = 7.29$, $p = 0.001$, $\eta 2 = 0.132$). Children updated their effort belief less after a failure compared to adolescents (children vs early adolescents: $t(63) = -2.77$, $p = 0.007$ $d = 0.708$ ; children vs late adolescents: $t(59) = -4.06$, $p < 0.001$, $d = 1.039$; early vs late adolescents: $t(70) = 0.83$, $p = 0.412$, $d = 0.196$). This shows that both adolescent groups used the information from their previous actions more to update beliefs, thus learning more than children. Additionally, all the groups took previous reward into account similarly ($F(2,96) = 0.18$, $p = 0.839$, $\eta 2 = 0.004$). They also used their own effort belief from previous trial to estimate their current belief similarly ($F(2,96) = 1.14$, $p = 0.323$, $\eta 2 = 0.023$), showing that all groups incorporated their own beliefs to construct effort beliefs on current trial.

*Effort learning for all groups*

To assess effort learning, we performed a multiple regression analysis that predicted how much physical effort was exerted on every trial with button presses. Replicating our previous results (Hauser, Eldar, & Dolan, 2017), we found that there was a significant effect of previous effort (Fig S5A, $t(98) = 8.82$, $p < 0.001$), failure ($t(98) = 10.22$, $p < 0.001$) and reward ($t(98) = 7.05$, $p < 0.001$). The effect of previous effort indicates that participants did not exert effort randomly but took into account their previous action. The previous failure effect implies that participants exerted effort according to their previous failure and learned from it. Subsequent analysis showed that participants compensated their failure by increasing effort after failure ($M =14.59$, $SD = 6.938$, $t(98) = 20.71$, $p < 0.001$) and decreasing it after a successful trial ($M = -4.66$, $SD = 3.002$, $t(98) = -15.44$, $p < 0.001$) – a clear sign that they dynamically adapted the effort based on previous performance (Hauser et al., 2017). The weight of the previous reward on the effort exerted highlights the motivation to apply more effort on high-value trials.

*Effort learning develops over childhood and adolescents*

We found that there were age effects on learning how much effort to exert. Firstly, there were age effect on how participants approximated their previous effort exertion to execute effort (Fig S5A; $F(2,96) = 7.34$, $p = 0.001$, $\eta 2 = 0.133$). Children did not use the information about how much effort they used in the previous trial as much as early adolescents ($t(63) = -2.23$, $p = 0.029$, $d = 0.556$) and late adolescents ($t(59) = -3.87$, $p < 0.001$, $d = 0.982$). There was also a trend of early adolescents taking previous effort less into account compared to late adolescents ($t(70) = 1.74$, $p = 0.087$, $d = 0.411$). Secondly, there were age effects on how the groups exerted effort after a failure ($F(2,96) = 5.06$, $p = 0.008$, $\eta 2 = 0.095$). Children had similar sensitivity to failure compared to early adolescents ($t(63) = -1.49$, $p = 0.143$, $d = 0.375$), but were less sensitive to failure compared to late adolescents $t(59) = -3.00$, $p = 0.004$, $d = 0.785$). There was marginal difference in failure sensitivity between early and late adolescents ($t(70) = 1.91$, $p = 0.060$, $d = 0.449$). Both of these effects together show that the learning abilities develop over time as it is very important to incorporate information about one's own behaviour (this case their exerted effort) and the environment's feedback (this case failure) for a good learning strategy. Previous reward affected the exerted effort similarly between all ages ($F(2,96) = 0.87$, $p = 0.421$, $\eta 2 = 0.018$), showing that all groups were more motivated by higher rewards, but did not use this information differently to exert effort.

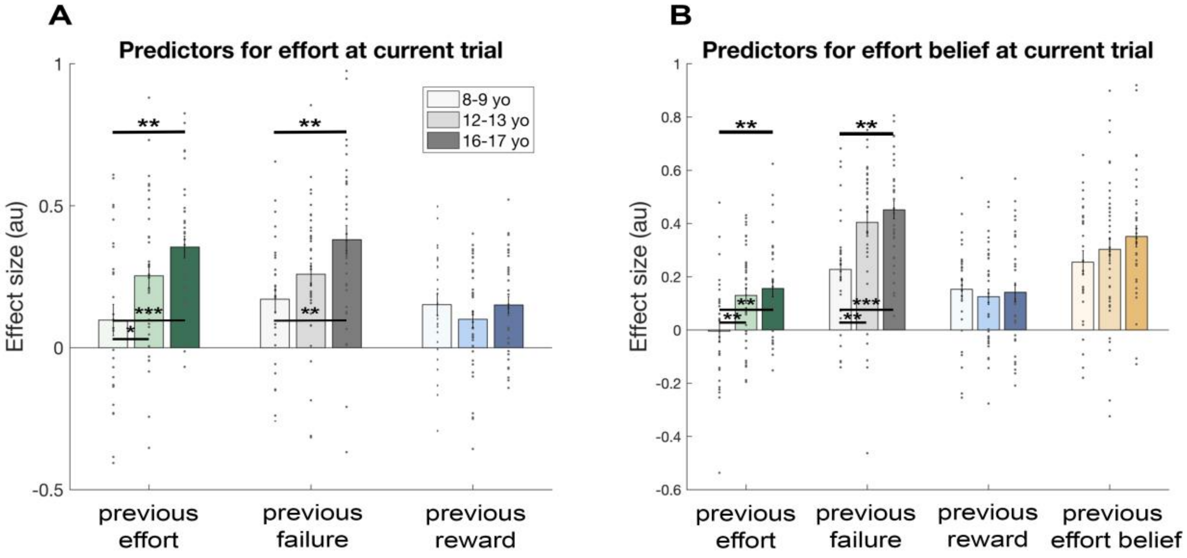


**Figure S5.** Effort learning behaviour (A) Analysis shows that the exerted effort was predicted by factors of the previous effort, failure to surpass the threshold on the previous trial, and the reward magnitude on previous trial, demonstrating that subjects successfully learned about reward and effort requirements. Additionally, the factor of previous effort and failure to surpass the threshold change over development. The factor of previous effort is lower in children compared to the adolescent groups, and the previous failure factor is lower in children compared to late adolescents (16-17 yo). (B) Analysis split by groups showing that effort belief was predicted by factors of previous effort, previous failure, reward and participant's own previous reported effort belief. Previous effort and previous failure factors are lower in predicting effort belief in children (8-9 year-old) compared to adolescents (12-13 yo and 16-17 yo). *** p<.001; ** p<.01; * p<.05; yo year-olds.

### References


Hauser, T. U., Eldar, E., & Dolan, R. J. (2017). Separate mesocortical and mesolimbic pathways encode effort and reward learning signals. *Proceedings of the National Academy of Sciences of the United States of America*. https://doi.org/10.1073/pnas.1705643114

Lefebvre, G., Lebreton, M., Meyniel, F., Bourgeois-Gironde, S., & Palminteri, S. (2017). Behavioural and neural characterization of optimistic reinforcement learning. *Nature Human Behaviour*. https://doi.org/10.1038/s41562-017-0067

Rescorla, R. A., & Wagner, A. R. (1972). *A theory of Pavlovian conditioning: variations in the effectiveness of reinforcement and nonreinforcement*. (A. H. Black & W. F. Prokasy, Eds.), *Classical Conditioning II: Current Research and Theory*. New York: Appleton-Century-Crofts.

Schwarz, G. (1978). Estimating the Dimension of a Model. *The Annals of Statistics*. https://doi.org/10.1214/aos/1176344136
